# BED-domain containing immune receptors confer diverse resistance spectra to yellow rust

**DOI:** 10.1101/299651

**Authors:** Clemence Marchal, Jianping Zhang, Peng Zhang, Paul Fenwick, Burkhard Steuernagel, Nikolai M. Adamski, Lesley Boyd, Robert McIntosh, Brande B.H. Wulff, Simon Berry, Evans Lagudah, Cristobal Uauy

## Introductory paragraph

Crop diseases reduce wheat yields by ~25% globally and thus pose a major threat to global food security^1^. Genetic resistance can reduce crop losses in the field and can be selected for through the use of molecular markers. However, genetic resistance often breaks down following changes in pathogen virulence, as experienced with the wheat yellow (stripe) rust fungus *Puccinia striiformis* f. sp. *tritici* (*Pst*)*^2^*. This highlights the need to (i) identify genes that alone or in combination provide broad-spectrum resistance and (ii) increase our understanding of the underlying molecular modes of action. Here we report the isolation and characterisation of three major yellow rust resistance genes (*Yr7, Yr5*, and *YrSP*) from hexaploid wheat (*Triticum aestivum*), each having a distinct and unique recognition specificity. We show that *Yr5*, which remains effective to a broad range of *Pst* isolates worldwide, is allelic to *YrSP* and paralogous to *Yr7*, both of which have been overcome by multiple *Pst* isolates. All three *Yr* genes belong to a complex resistance gene cluster on chromosome 2B encoding nucleotide-binding and leucine-rich repeat proteins (NLRs) with a non-canonical N-terminal zinc-finger BED domain^3^ that is distinct from those found in non-NLR wheat proteins. We developed diagnostic markers to accelerate haplotype analysis and for marker-assisted selection to enable the stacking of the non-allelic *Yr* genes. Our results provide evidence that the BED-NLR gene architecture can provide effective field-based resistance to important fungal diseases such as wheat yellow rust.

## Main

In plant immunity, NLRs act as intracellular immune receptors that upon pathogen recognition trigger a series of signalling steps that ultimately lead to cell death, thus preventing the spread of infection^4,5^. The NB-ARC domain is the hallmark of NLRs which in most cases include leucine-rich repeats (LRRs) at the C-terminus. Recent *in silico* analyses have identified NLRs with additional ‘integrated’ domains^6–8^, including zinc-finger BED domains (BED-NLRs). The BED domain function within BED-NLRs is unknown, although the BED domain from the non-NLR DAYSLEEPER protein was shown to bind DNA in *Arabidopsis*^9^. BED-NLRs are widespread across Angiosperm genomes^6–8^ and this gene architecture has been shown to confer resistance to bacterial blast in rice (*Xa1*^10,11^).

The genetic relationship between *Yr5* and *Yr7* has been debated for almost 45 years^12,13^. Both genes map to chromosome arm 2BL in hexaploid wheat and were hypothesized to be allelic^14^, and closely linked with *YrSP*^15^. Whilst only one of >6,000 tested *Pst* isolates worldwide has been found virulent to *Yr5* (Supplementary Table 1^14,16^), both *Yr7* and *YrSP* have been overcome in the field. For *Yr7*, this is likely due to its wide deployment in cultivars (Supplementary Table 2, Supplementary Figure 1). This highlights the importance of stewardship plans (including diagnostic markers) to deploy *Yr5* in combination with other genes as currently done in the US (e.g. *Yr5+Yr15;* UC Davis breeding programme).

To clone the genes encoding *Yr7, Yr5*, and *YrSP*, we identified susceptible ethyl methanesulfonate-derived (EMS) mutants from different genetic backgrounds carrying these genes (Figure 1, Supplementary Tables 3-4). We performed MutRenSeq^17^ and isolated a single candidate contig for each of the three genes based on nine, ten, and four independent susceptible mutants, respectively (Figure 1a; Supplementary Figure 2). The three candidate contigs were genetically linked to a common mapping interval, previously identified for the three *Yr* loci^15,18,19^. Their closest homologs in the Chinese Spring wheat genome sequence (RefSeq v1.0) all lie within this common genetic interval (Figure 1b; Supplementary Figure 3).

**Figure 1:**
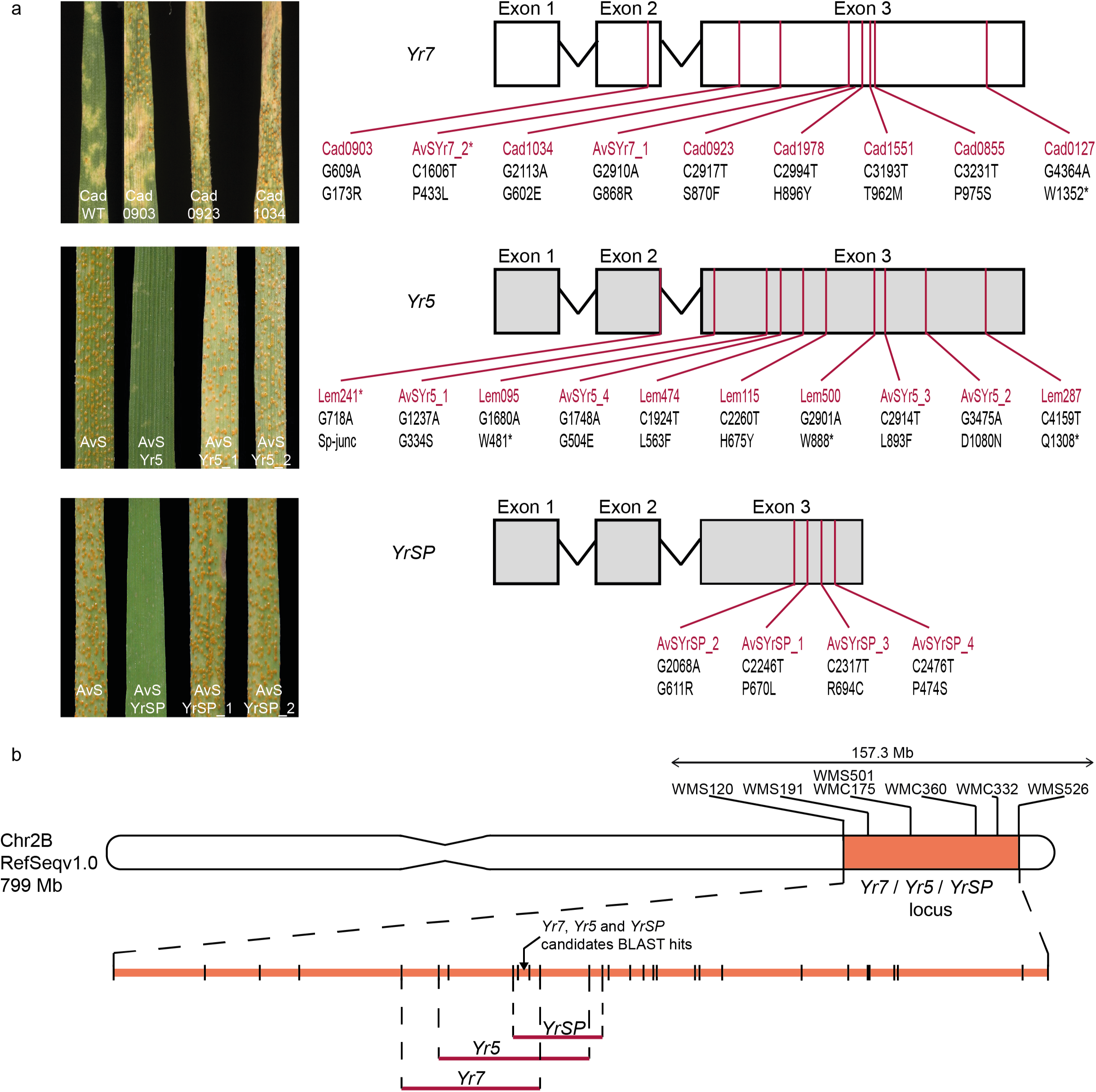
*Yr5* and *YrSP* are allelic, but paralogous to *Yr7*. **a**, Left: Wild-type and selected EMS-derived susceptible mutant lines for *Yr7, Yr5*, and *YrSP* (Supplementary Table 3 and 3) inoculated with *Pst* isolate 08/21 (*Yr7*)*, Pst* 150 E16 A+ (*Yr5*), or *Pst* 134 E16 A+ (*YrSP*). Right: Candidate gene structures, with mutations in red, and their predicted effects on the translated protein. **b**, Schematic representation of the physical interval of the *Yr* loci. The *Yr7/Yr5/YrSP* locus is shown in orange on chromomsome 2B with previously published SSR markers in black. Markers developed in this study to confirm the genetic linkage between the phenotype and the candidate contigs are shown as black lines underneath the chromosme. *Yr* loci mapping intervals are defined by the red horizontal lines. A more detailed genetic map is shown in Supplementary Figure 3.

Within each contig we predicted a single open reading frame based on RNA-Seq data. All three predicted *Yr* genes displayed similar exon-intron structures (Figure 1a), although *YrSP* was truncated in exon 3 due to a single base deletion that resulted in a premature termination codon. The DNA sequences of *Yr7* and *Yr5* were 77.9% identical across the complete gene; whereas *YrSP* was a truncated version of *Yr5*, sharing 99.8% identity in the common sequence (Supplementary Files 1 and 2). This suggests that *Yr5* and *YrSP* are encoded by alleles of the same gene, but are paralogous to *Yr7*. The 23 mutations identified by MutRenSeq were confirmed by Sanger sequencing and all lead to either an amino acid substitution or a truncation allele (splice junction or termination codon) (Figure 1a; Supplementary Table 4). Taken together, the mutant and genetic analyses demonstrate that *Yr5* and *YrSP* are allelic, while *Yr7* is encoded by a related, yet distinct gene.

The Yr7, Yr5, and YrSP proteins contain a zinc-finger BED domain at the N-terminus, followed by the canonical NB-ARC domain. Unlike previously cloned resistance genes in grasses (e.g. *Mla10l*^20^, *Sr33*^21^, *Pm*3^22^), neither *Yr7* nor *Yr5/YrSP* encode Coiled Coil domains at the N-terminus (Supplementary Figure 4). Only Yr7 and Yr5 proteins encode multiple LRR motifs at the C-terminus (Figure 2a; green bars), YrSP having lost most of the LRR region due to the presence of the premature termination codon in exon 3. YrSP still confers functional resistance to *Pst*, although with a different recognition specificity to Yr5. Yr7 and Yr5/YrSP are highly conserved in the N-terminus, with a single amino-acid change in the BED domain. This high degree of conservation is eroded downstream of the BED domain (Figure 2a). The BED domain is required for Yr7-mediated resistance, as a single amino acid change in mutant line Cad0903 led to a susceptible reaction (Figure 1a). However, recognition specificity is not solely governed by the BED domain, as the *Yr5* and *YrSP* alleles have identical BED domain sequences, yet confer resistance to different *Pst* isolates. The highly conserved Yr7 and Yr5/YrSP BED domains could function in a similar way to the integrated WKRY domain in the *Arabidopsis* RRS1-R immune receptor which binds unrelated bacterial effectors yet activates defense response through mechanisms involving other regions of the protein^23^.

**Figure 2:**
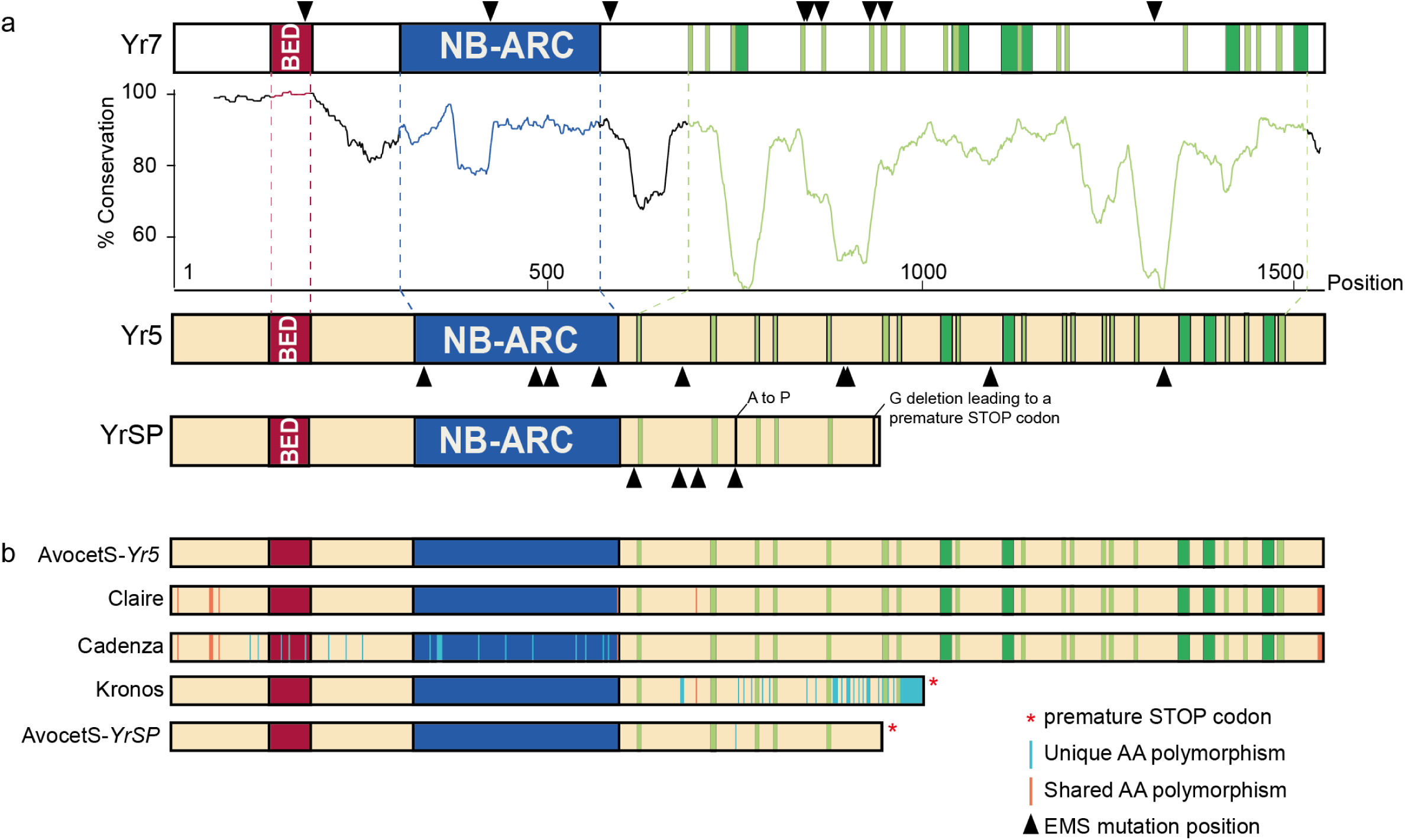
*Yr7* and *Yr5/YrSP* encode integrated BED-domain immune receptor genes. **a**, Schematic representation of the Yr7, Yr5, and YrSP protein domain organisation. BED domains are highlighted in red, NB-ARC domains are in blue, LRR motifs from NLR-Annotator are in dark green, and manually annotated LRR motifs (xxLxLxx) are in light green. Black triangles represent the EMS-induced mutations within the protein sequence. The plot shows the degree of amino acid conservation (50 amino acid rolling average) between Yr7 and Yr5 proteins, based on the conservation diagram produced by Jalview (2.10.1) from the protein alignment. Regions that correspond to the conserved domains have matching colours. The amino acid changes between Yr5 and YrSP are annotated in black on the YrSP protein. **b**, Five Yr5/YrSP haplotypes were identified in this study. Polymorphisms are highlighted across the protein sequence with orange vertical bars for polymorphisms shared by at least two haplotypes and blue vertical bars for polymorphisms that are unique to the corresponding haplotype. Matching colours across protein structures illustrate 100% sequence conservation.

We examined the allelic variation in *Yr7, Yr5*, and *YrSP* across eight sequenced tetraploid and hexaploid wheat genomes (Supplementary Table 5). We identified *Yr7* only in Cadenza and Paragon, which are identical-by-descent in this interval (Supplementary File 3, Supplementary Table 6, and Supplementary Figure 5). Both cultivars are derived from the original source of *Yr7*, tetraploid durum wheat (*T. turgidum* ssp. *durum*) cultivar Iumillo and its hexaploid derivative Thatcher (Supplementary Figure 5). None of the three sequenced tetraploid accessions (Svevo, Kronos, Zavitan) carry *Yr7* (Supplementary Table 6).

For *Yr5/YrSP*, we identified three additional alleles in the sequenced hexaploid wheat cultivars (Figure 2b; Supplementary Table 7). Cultivar Claire encodes a complete NLR with six amino-acid changes, including one within the NB-ARC domain, and six polymorphisms in the C-terminus compared to Yr5. Cultivars Robigus, Paragon, and Cadenza also encode a full length NLR that shares common polymorphisms with Claire, in addition to 19 amino acid substitutions across the BED and NB-ARC domains. The C-terminus polymorphisms between Yr5 and the other cultivars is due to a 774 bp insertion in *Yr5*, close to the 3’ end, which carries an alternate termination codon (Supplementary File 2). Tetraploid cultivars Kronos and Svevo encode a fifth Yr5/YrSP allele with a truncation in the LRR region distinct from YrSP, in addition to multiple amino acid substitutions across the C-terminus (Supplementary Table 7). This truncated tetraploid allele is reminiscent of YrSP and is expressed in Kronos (see Methods). However, none of these cultivars (Claire, Robigus, Paragon, Cadenza, Svevo, and Kronos) exhibit a *Yr5/YrSP* resistance response, suggesting that these amino acid changes and truncations may alter recognition specificity or protein function.

We designed diagnostic markers for *Yr7, Yr5*, and *YrSP* to facilitate their detection and use in breeding. We confirmed their presence in the donor cultivars Thatcher and Lee (*Yr7*), Spaldings Prolific (*YrSP*), and spelt wheat cv. Album (*Yr5*) (Supplementary Tables 8-9; Supplementary Figures 5-6). We tested *Yr7* and *YrSP* markers in a collection of global landraces^24^ and European cultivars^25^ released over the past one hundred years. *YrSP* was absent from the tested germplasm, except for AvocetS-*YrSP* (Supplementary Table 9). *Yr7* on the otherhand was more prevalent in the germplasm tested and we could track its presence across pedigrees, including Cadenza derived cultivars (Supplementary Tables 8-9; Supplementary Figure 5). We confirmed *Yr5* in the AvocetS-Yr5 and Lemhi-*Yr5* lines and it was not detected in the other tested lines, consistent with the fact that *Yr5* has not yet been deployed within European breeding programmes (Supplementary Tables 10 and 17 and Supplementary Figure 6). The *Yr5* diagnostic marker will facilitate its deployment, hopefully within a breeding strategy that ensures its effectiveness long-term^26^.

We defined the *Yr7/Yr5/YrSP* syntenic interval across the wheat genomes and related grass species *Aegilops tauschii* (D genome progenitor), *Hordeum vulgare* (barley), *Brachypodium distachyon*, and *Oryza sativa* (rice) (Supplementary files 4 and 5, Supplementary Figure 7). We identified both canonical NLRs, as well as BED-NLRs across all genomes and species, except for barley, which only contained canonical NLRs across the syntenic region. The phylogenetic relationship based on the NB-ARC domain suggests a common evolutionary origin of these integrated domain NLR proteins before the wheat-rice divergence (~50 Mya) and an expansion in the number of NLRs in the A and B genomes of polyploid wheat species (Figure 3a; Supplementary Figure 8). Within the interval we also identified several genes in the A, B, and D genomes that encode two consecutive in-frame BED domains (named BED-I and BED_II; Figure 3b-c, Supplementary Figure 7) followed by the canonical NLR. The BED domains in these genes were fully encoded within a single exon (exons 2 and 3) and in most cases had a four-exon structure (Figure 3c). This is consistent with the three-exon structure of single BED domain genes, such as *Yr7* and *Yr5/YrSP* (BED-I encoded on exon 2). To our knowledge this is the first report of the double BED domain NLR protein structure. The biological function of this molecular innovation remains to be determined, although our data show that the single BED-I structure can confer *Pst* resistance and is required for *Yr7*-mediated resistance.

**Figure 3:**
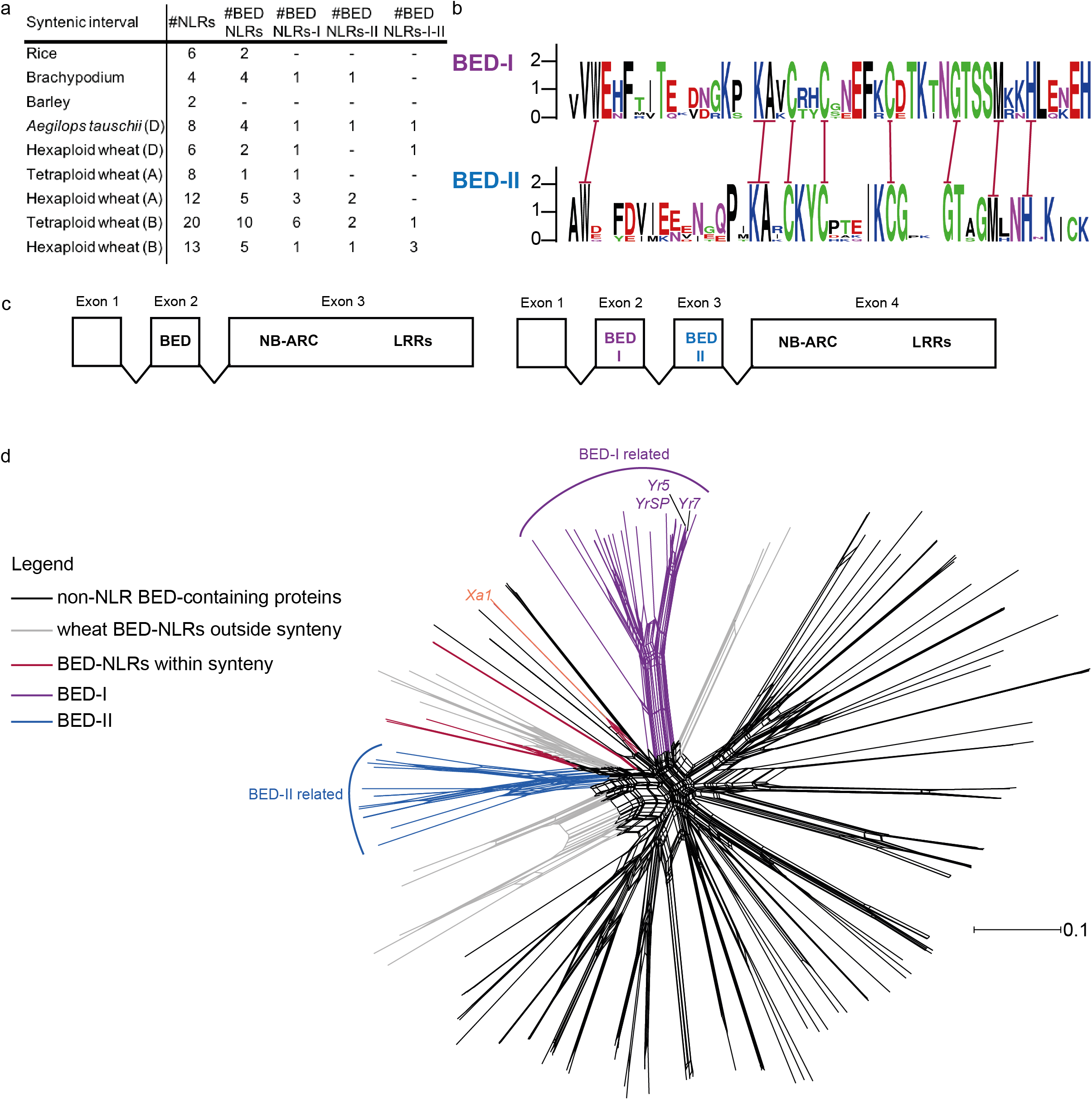
BED domains from BED-NLRs and non-NLR proteins are distinct. **a**, Numbers of NLRs in the syntenic regions across grass genomes (see Supplementary Figure 7), including BED-NLRs. **b**, WebLogo (http://weblogo.berkeley.edu/logo.cgi) diagram showing that the BED-I and BED-II domains are distinct, with only the highly conserved residues that define the BED domain (red bars) being conserved between the two types. **c**, Gene structure most commonly observed for BED-NLRs and BED-BED-NLRs within the *Yr7/Yr5/YrSP* syntenic interval. **d**, Neighbour-net analysis based on uncorrected *P* distances obtained from alignment of 153 BED domains including the 108 BED-containing proteins (including 25 NLRs) from RefSeq v1.0, BED domains from NLRs located in the syntenic region as defined in Supplementary Figure 7, and BED domains from Xa1 and ZBED from rice. BED-I and II clades are highlighted in purple and blue, respectively. BED domains from the syntenic regions not related to either of these types are in red. BED domains derived from non-NLR proteins are in black and BED domains from BED-NLRs outside the syntenic region are in grey. Seven BED domains from non-NLR proteins were close to BED domains from BED-NLRs. Supplementary Figure 9 includes individual labels.

Among other mechanisms, integrated domains of NLRs are hypothesised to act as decoys for pathogen effector targets^5^. This would suggest that the integrated domain might be sequence-related to the host protein targeted by the effector. To identify these potential effector targets in the host, we retrieved all BED-domain proteins (108) from the hexaploid wheat genome, including 25 BED-NLRs, and additional BED-NLRs located in the syntenic intervals (Supplementary Table 11; Supplementary file 4). We also retrieved the rice Xa1^10,11^ and ZBED proteins, the latter being hypothesized to mediate rice resistance to *Magnaporthe oryzae*^7^. We used the split network method implemented in SplitsTree4^27^ to represent the relationships between these BED domains (Figure 3d; Supplementary Figure 9). Overall, BED domains are diverse, although there is evidence of a split between BED domains from BED-NLRs and non-NLR proteins (only 7 of 83 non-NLRs clustered with the BED-NLRs). Given that the base of the split is broad, integrated BED-domains most likely derive from multiple integration events. However, Yr7 and Yr5/YrSP both arose from a common integration event that occurred before the *Brachypodium*-wheat divergence (Supplementary Figure 9, purple). This is consistent with the hypothesis that integrated domains might have evolved to strengthen the interaction with pathogen effectors after integration^28^, although we cannot exclude the potential role of the BED domains in signalling at this stage.

Among BED-NLRs, BED-I and BED-II constitute two major clades, consistent with their relatively low amino acid conservation (Figure 3b), that are comprised solely of genes from within the *Yr7/Yr5/YrSP* syntenic region. Seven non-NLR BED domain wheat proteins clustered with BED-NLRs. These are most closely related to the *Brachypodium* and rice BED-NLR proteins and were not expressed in RNA-Seq data from a *Yr5* time-course (re-analysis of published data^29^; Supplementary Figure 10, Supplementary Table 12). Similarly, no BED-containing protein was differentially expressed during this infection time-course, consistent with the prediction that effectors alter their targets’ activity at the protein level in the integrated-decoy model^5^. We cannot however disprove that these closely related BED-containing proteins are involved in BED-NLR-mediated resistance.

BED-NLRs are frequent in Triticeae, and occur in other monocot and dicot tribes^6–8^. To date a single BED-NLR gene, *Xa1*, has been shown to confer resistance to plant pathogens^10,11^. In the present study, we show that the distinct *Yr7, Yr5*, and *YrSP* resistance specificities belong to a complex NLR cluster on chromosome 2B and are encoded by two paralogous BED-NLRs genes. We report an allelic series for the *Yr5/YrSP* gene with five independent alleles, including three full-length BED-NLRs (including *Yr5*) and two truncated versions (including *YrSP*). This wider allelic series could be of functional significance as previously shown for the *Mla* and *Pm3* loci that confer resistance to *Blumeria graminis*^22,30^ in barley and wheat, respectively, and the flax *L* locus conferring resistance to *Melampsora lini^31^*. Overall, our results add strong evidence for the importance of the BED-NLR architecture in plant-pathogen interactions. The paralogous and allelic relationship of these three distinct *Yr* loci will inform future hypothesis-driven engineering of novel recognition specificities.

## Methods

### MutRenSeq

#### Mutant identification

Supplementary Table 3 summarises plant materials and *Pst* isolates used to identify mutants for each *Yr* gene. We used an EMS-mutagenised population in cultivar Cadenza^32^ to identify mutants in *Yr7;* whereas EMS-populations in the corresponding AvocetS-*Yr* near isogenic line (NIL) were used to identify *Yr5* and *YrSP* mutants. For *Yr7*, we inoculated M_3_ plants from the Cadenza EMS population with *Pst* isolate 08/21 which is virulent to *Yr1, Yr2, Yr3, Yr4, Yr6, Yr9, Yr17, Yr27, Yr32, YrRob*, and *YrSol*^33^. We hypothesised that susceptible mutants would carry mutations in *Yr7*. Plants were grown in 192-well trays in a confined glasshouse with no supplementary lights or heat. Inoculations were performed at the one leaf stage (Zadoks 11) with a talc – urediniospore mixture. Trays were kept in darkness at 10 °C and 100% humidity for 24 hours. Infection types (IT) were recorded 21 days post-inoculation (dpi) following the Grassner and Straib scale^34^. Identified susceptible lines were progeny tested to confirm the reliability of the phenotype and DNA from M4 plants was used for RenSeq (see section below). Similar methods were used for AvocetS-*Yr7*, AvocetS-*Yr5*, and AvocetS-*YrSP* EMS-mutagenised populations with the following exceptions: *Pst* pathotypes 108 E141 A+ (University of Sydney Plant Breeding Institute Culture no. 420), 150 E16 A+ (Culture no. 598) and 134 E16 A+ (Culture no. 572) were used to evaluate *Yr7, Yr5*, and *YrSP* mutants, respectively. EMS-derived susceptible mutants in Lemhi-*Yr5* were previously identified^35^ and DNA from M5 plants was used for RenSeq.

#### DNA preparation, resistance gene enrichment and sequencing (RenSeq)

We extracted total genomic DNA from young leaf tissue using the large-scale DNA extraction protocol from the McCouch Lab (https://ricelab.plbr.cornell.edu/dna_extraction) and a previously described method^36^. We checked DNA quality and quantity on a 0.8% agarose gel and with a NanoDrop spectrophotometer (Thermo Scientific). Arbor Biosciences (Ann Arbor, MI, USA) performed the targeted enrichment of NLRs according to the MYbaits protocol using an improved version of the previously published Triticeae bait library available at github.com/steuernb/MutantHunter. Library construction was performed using the TruSeq RNA protocol v2 (Illumina 15026495). Libraries were pooled with one pool of samples for Cadenza mutants and one pool of eight samples for the Lemhi-*Yr5* parent and Lemhi-*Yr5* mutants. AvocetS-Yr5 and AvocetS-*YrSP* wild-type, together with their respective mutants, were also processed according to the MYbaits protocol and the same bait library was used. All enriched libraries were sequenced on a HiSeq 2500 (Illumina) in High Output mode using 250 bp paired end reads and SBS chemistry. For the Cadenza wild-type, we generated data on an Illumina MiSeq instrument. In addition to the mutants, we also generated RenSeq data for Kronos and Paragon to assess the presence of *Yr5* in Kronos and *Yr7* in Paragon. Details of all the lines sequenced, alongside NCBI accession numbers, are presented in Supplementary Tables 3 and 12.

### MutantHunter pipeline

We adapted the pipeline from https://github.com/steuernb/MutantHunter/ to identify candidate contigs for the targeted *Yr* genes. First, we trimmed the RenSeq-derived reads with trimmomatic^37^ using the following parameters: ILLUMINACLIP:TruSeq2-PE.fa:2:30:10 LEADING:30 TRAILING:30 SLIDINGWINDOW:10:20 MINLEN:50 (v0.33). We made *de novo* assemblies of wild-type plant trimmed reads with the CLC assembly cell and default parameters apart from the word size (-w) parameter that we set to 64 (v5.0, http://www.clcbio.com/products/clc-assembly-cell/) (Supplementary Table 14). We then followed the MutantHunter pipeline detailed at https://github.com/steuernb/MutantHunter/. For Cadenza mutants, we used the following MutantHunter program parameters to identify candidate contigs: -c 20 -n 6 -z 1000. These options require a minimum coverage of 20x for SNPs to be called; at least six susceptible mutants must have a mutation in the same contig to report it as candidate; small deletions were filtered out by setting the number of coherent positions with zero coverage to call a deletion mutant at 1000. The -n parameter was modified accordingly in subsequent runs with the Lemhi-*Yr5* datasets (-n 6).

To identify *Yr5* and *YrSP* contigs from Avocet mutants, we followed the MutantHunter pipeline with all default parameters, except in the use of CLC Genomics Workbench (v10) for reads QC, trimming, *de novo* assembly of Avocet wild-type and mapping all the reads against *de novo* wild-type assembly. The default MutantHunter parameters were used except that −z was set as 100. The parameter −n was set to 2 in the first run and then to 3 in the second run. Two *Yr5* mutants were most likely sibling lines as they carried identical mutations at the same position (Supplementary Figure 2, Supplementary Table 4).

For *Yr7* we identified a single contig with six mutations, however we did not identify mutations in line Cad0903. Upon examination of the *Yr7* candidate contig we predicted that the 5’ region was likely to be missing (Supplementary Figure 2). We thus annotated potential NLRs in the Cadenza genome assembly available from the Earlham Institute (Supplementary Table 5, http://opendata.earlham.ac.uk/Triticum_aestivum/EI/v1.1) with the NLR-Annotator program using default parameters (https://github.com/steuernb/NLR-Annotator). We identified an annotated NLR in the Cadenza genome with 100% sequence identity to the *Yr7* candidate contig, which extended beyond our *de novo* assembled sequence. We therefore replaced the previous candidate contig with the extended Cadenza sequence (100% sequence identity) and mapped the RenSeq reads from Cadenza wild-type and mutants as described above. This confirmed the candidate contig for *Yr7* as we retrieved the missing 5’ region including the BED domain. The improved contig now also contained a mutation in the outstanding mutant line Cad0903 (Supplementary Figure 2). The Triticeae bait library does not include integrated domains in its design so they are prone to be missed, especially when located at the ends of an NLR. Sequencing technology could also have accounted for this: MiSeq was used for Cadenza wild-type whereas HiSeq was chosen for Lemhi-*Yr5* and we recovered the 5’ region in the latter, although coverage was lower than for the regions encoding canonical domains. In summary, we sequenced nine, ten, and four mutants for *Yr7, Yr5*, and *YrSP*, respectively and identified for each target gene a single contig that accounted for all mutants.

### Candidate contig confirmation and gene annotation

We sequenced the *Yr7, Yr5*, and *YrSP* candidate contigs from the mutant lines (annotated in Supplementary Files 1 and 2) to confirm the EMS-derived mutations using primers documented in Supplementary Table 15. We first PCR-amplified the complete locus from the same DNA preparations as the ones submitted for RenSeq with the Phusion^®^ High-Fidelity DNA Polymerase (New England Biolabs) following the suppliers protocol (https://www.neb.com/protocols/0001/01/01/pcr-protocol-m0530). We then carried out nested PCR on the obtained product to generate overlapping 600-1,000 bp amplicons that were purified using the MiniElute kit (Qiagen). The purified PCR products were sequenced by GATC following the LightRun protocol (https://www.gatc-biotech.com/shop/en/lightrun-tube-barcode.html). Resulting sequences were aligned to the wild-type contig using ClustalOmega (https://www.ebi.ac.uk/Tools/msa/clustalo/). This allowed us to curate the *Yr7* locus in the Cadenza assembly that contained two sets of unknown (‘N’) bases in its sequence, corresponding to a 39 bp insertion and a 129 bp deletion (Supplementary File 3), and to confirm the presence of the mutations in each mutant line.

We used HISATt2^38^ (v2.1) to map RNA-Seq reads available from Cadenza and AvocetS-*Yr5*^29^ to the RenSeq *de novo* assemblies with curated loci to define the structure of the genes. We used the following parameters: --no-mixed --no-discordant to map reads in pairs only. We used the --novel-splicesite-outfile to predict splicing sites that we manually scrutinised with the genome visualisation tool IGV^39^ (v2.3.79). Predicted coding sequences (CDS) were translated using the ExPASy online tool (https://web.expasy.org/translate/). This allowed us to predict the effect of the mutations on each candidate transcript (Figure 1a; Supplementary Table 4). The long-range primers for both *Yr7* and *Yr5* loci were then used on the corresponding susceptible Avocet NIL mutants to determine whether the genes were present and carried mutations in that background (Figure 1a; Supplementary Files 1 and 2).

### Coiled coil domain prediction

To determine whether *Yr7, Yr5*, and *YrSP* encode Coiled Coil (CC) domains we used the NCOILS prediction program^40^ (v1.0, https://embnet.vital-it.ch/software/COILS_form.html) with the following parameters: MTIK matrix with applying a 2.5-fold weighting of positions a,d. We compared the profiles to those obtained with already characterised CC-NLR encoding genes *Sr33, Mla10, Pm3* and *RPS5* (Supplementary Figure 4). We also ran the program on Yr7 and Yr5 protein sequences where the BED domain was manually removed to determine whether its integration could have disrupted an existing CC domain. To further investigate whether *Yr7*, *Yr5*, and *YrSP* encode CC domains we performed a BLASTP analysis^41^ with their N-terminal region, from the methionine to the first amino acid encoding the NB-ARC domain, with or without the BED domain (Supplementary Figure 4).

### Genetic linkage

We generated a set of F2 populations to genetically map the candidate contigs (Supplementary Table 3). For *Yr7* we developed an F_2_ population based on a cross between the susceptible mutant line Cad0127 to the Cadenza wild-type (population size 139 individuals). For *Yr5* and *YrSP* we developed F2 populations between AvocetS and the NILs carrying the corresponding *Yr* gene (94 individuals for *YrSP* and 376 for *Yr5*). We extracted DNA from leaf tissue at the seedling stage (Zadoks 11) following a previously published protocol^42^ and Kompetitive Allele Specific PCR (KASP) assays were carried out as described in^43^. R/qtl package^44^ was used to produce the genetic map based on a general likelihood ratio test and genetic distances were calculated from recombination frequencies (v1.41-6).

We used previously published markers linked to *Yr7, Yr5*, and *YrSP* (WMS526, WMS501 and WMC175, WMC332, respectively^15,18,19^) in addition to closely linked markers WMS120, WMS191, and WMC360 (based on the GrainGenes database https://wheat.pw.usda.gov/GG3/) to define the physical region on the Chinese Spring assembly RefSeq v1.0 (https://wheat-urgi.versailles.inra.fr/Seq-Repository/Assemblies). Two different approaches were used for genetic mapping depending on the material. For *Yr7*, we used the public data^32^ for Cad0127 (www.wheat-tilling.com) to identify nine mutations located within the *Yr7* physical interval based on BLAST analysis against RefSeq v1.0. We used KASP primers when available and manually designed additional ones including an assay targeting the Cad0127 mutation in the *Yr7* candidate contig (Supplementary Table 15). We genotyped the Cad0127 F_2_ populations using these nine KASP assays and confirmed genetic linkage between the Cad0127 *Yr7* candidate mutation and the nine mutations across the physical interval (Supplementary Figure 3).

For *Yr5* and *YrSP*, we first aligned the candidate contigs to the best BLAST hit in an AvocetS RenSeq *de novo* assembly. We then designed KASP primers targeting polymorphisms between these sequences and used them to genotype the corresponding F_2_ population (Supplementary Table 15). For both candidate contigs we confirmed genetic linkage with the previously published genetic intervals for these *Yr* genes (Supplementary Figure 3).

### *Yr7* gene-specific markers

We aligned the *Yr7* sequence with the best BLAST hits in the genomes listed on Supplementary Table 5 and designed KASP primers targeting polymorphisms that were *Yr7-*specific. Three markers were retained after testing on a selected panel of Cadenza-derivatives and cultivars that were positive for *Yr7* markers in the literature, including the *Yr7* reference cultivar Lee (Supplementary Table 8, 8 and 15). The panel of Cadenza-derivatives was phenotyped with three *Pst* isolates: *Pst* 08/21 (Yr7-avirulent), *Pst* 15/151 (Yr7-avirulent – virulent to *Yr1, 2, 3, 4, 6, 9, 17, 25, 32, Rendezvous, Sp, Robigus, Solstice*) and *Pst* 14/106 (*Yr7-*virulent, virulent to *Yr1, 2, 3, 4, 6, 7, 9, 17, 25, 32, Sp, Robigus, Solstice, Warrior, Ambition, Cadenza, KWS Sterling, Apache*) to determine whether Yr7-positive cultivars, as identified by the three KASP markers, displayed a consistent specificity (Supplementary Table 8). Pathology assays were performed as for the screening of the Cadenza mutant population. We retrieved pedigree information for the analysed cultivars from the Genetic Resources Information System for Wheat and Triticale database (GRIS, www.wheatpedigree.net) and used the Helium software^45^ (v1.17) to illustrate the breeding history of *Yr7* in the UK (Supplementary Figure 5).

We used the three *Yr7* KASP markers to genotype (i) cultivars from the AHDB Wheat Recommended List from 2005-2018 (https://cereals.ahdb.org.uk/varieties/ahdb-recommended-lists.aspx); (ii) the Gediflux collection of European bread wheat cultivars released between 1920 and 2010^25^ and (iii) the core Watkins collection, which represents a global set of wheat landraces collected in the 1930s^24^. Results are reported in Supplementary Table 9.

### *Yr5* and *YrSP* gene-specific markers

We identified a 774 bp insertion in the *Yr5* allele 29 bp upstream of the STOP codon with respect to the Cadenza and Claire alleles. Genomic DNA from *YrSP* confirmed that the insertion was specific to *Yr5*. We used this polymorphism to design primers flanking the insertion and tested them on a subset of the collections mentioned above. We added 32 DNA sample from diverse accessions of *Triticum dicoccoides*, the wild progenitor of domesticated wheat (passport data shown in Supplementary Table 17). We included DNA from *Triticum aestivum* ssp. *spelta* var. album^35^ (*Yr5* donor) and Spaldings Prolific (*YrSP* donor) to assess their amplification profiles. PCR amplification was conducted using a touchdown programme: 10 cycles, −0.5 °C per cycle starting from 67 °C and the remaining 25 cycles at 62 °C. This allowed us to increase the specificity of the reaction. We observed three different profiles on the tested varieties; (i) a 1,281 bp amplicon in *Yr5* positive cultivars, (ii) a 507 bp amplicon in the alternate *Yr5* allele carriers, including AvocetS-*YrSP*, Cadenza, and Claire, and (iii) no amplification in other varieties. We sequenced the different amplicons and confirmed the insertion in *Yr5* compared to the alternate alleles (Supplementary File 2). The lack of amplicons in some varieties most likely respresents the absence of the loci in the tested varieties. For *YrSP*, we aligned the *YrSP* and *Yr5* sequences to design KASP primers targeting the G to C SNP between the two alleles (Supplementary File 2, Supplementary Table 16). We tested the marker by genotyping selected cultivars as controls and cultivars from the AHDB Wheat Recommended List from 2005-2018 (Supplementary Table 9).

### *In silico* allele mining for *Yr7* and *Yr5*

We used the *Yr7* and *Yr5* sequences to retrieve the best BLAST hits in the *T. aestivum* and *T. turgdium* wheat genomes listed in Supplementary Table 5. The best *Yr5* hits shared between 93.6 and 99.3% sequence identity, which was comparable to what was observed for alleles derived from the wheat *Pm3* (>97% identity)^46^ and flax *L* (>90% identity)^31^ genes. *Yr7* was identified only in Paragon and Cadenza (Supplementary Table 6; See Supplementary File 3 for curation of the Paragon sequence).

### Analysis of the *Yr7* and *Yr5/YrSP* cluster on RefSeq v1.0

#### Definition of syntenic regions across grass genomes

We used NLR-Annotator to identify putative NLR loci on RefSeq v1.0 chromosome 2B and identified the best BLAST hits to *Yr7* and *Yr5* on RefSeq v1.0. Additional BED-NLRs and canonical NLRs were annotated in close physical proximity to these best BLAST hits. Therefore, to better define the NLR cluster we selected ten non-NLR genes located both distal and proximal to the region, and identified orthologs in barley, *Brachypodium*, and rice in *EnsemblPlants* (https://plants.ensembl.org/). We used different % ID cutoffs for each species (>92% for barley, >84% for *Brachypodium*, and >76% for rice) and determined the syntenic region when at least three consecutive orthologues were found. A similar approach was conducted for *Triticum* ssp and *Ae. tauschii* (Supplementary file 4).

#### Definition of the NLR content of the syntenic region

We extracted the previously defined syntenic region from the grass genomes listed in Supplementary Table 5 and annotated NLR loci with NLR-Annotator. We maintained previously defined gene models where possible, but also defined new gene models that were further analysed through a BLASTx analysis to confirm the NLR domains (Supplementary Files 4 and 5). The presence of BED domains in these NLRs was also confirmed by CD-Search (https://www.ncbi.nlm.nih.gov/Structure/cdd/wrpsb.cgi).

### Phylogenetic and neighbour network analyses

We aligned the translated NB-ARC domains from the NLR-Annotator output with MUSCLE^47^ using default parameters (v.3.8.31). We verified and manually curated the alignment with Jalview^48^ (v2.10.1). We used Gblocks^49^ (v0.91b) with the following parameters: Minimum Number Of Sequences For A Conserved Position: 9; Minimum Number Of Sequences For A Flanking Position: 14; Maximum Number Of Contiguous Nonconserved Positions: 8; Minimum Length Of A Block: 10; Allowed Gap Positions: None; Use Similarity Matrices: Yes; to eliminate poorly aligned positions. This resulted in 36% of the original 156 positions being taken forward for the phylogeny. We built a Maximum Likelihood tree with the RAxML^50^ program and the following parameters: raxmlHPC -f a -x 12345 -p 12345 -N 1000 -m PROTCATJTT -s <input_alignment.fasta> (MPI version v8.2.10). The best scoring tree with associated bootstrap values was visualised and mid-rooted with Dendroscope^51^ (v3.5.9). There was clear separation between NLRs belonging to the two different clusters but the sub-clades had less support. One explanation would be that conflicting phylogenetic signals that are due to events such as hybridization, horizontal gene transfer, recombination, or gene duplication and loss might have occurred in the region. Split networks allow nodes that do not represent ancestral species and can thus represent such incompatible and ambiguous signals. We therefore used this method in the following part of the analysis to analyse the relationship between the BED domains.

We used the Neighbour-net method^52^ implemented in SplitsTree4^27^ (v4.16) to analyse the relationships between BED domains from NLR and non-NLR proteins. First we retrieved all BED-containing proteins from RefSeq v1.0 using the following steps: we used hmmer (v3.1b2, http://hmmer.org/) to identify conserved domains in protein sequences from RefSeq v1.0. We applied a cut-off of 0.01 on i-evalue to filter out any irrelevant identified domains. We separated the set between NLR and non-NLRs based on the presence of the NB-ARC and sequence homology for single BED proteins. BED domains were extracted from the corresponding protein sequences based on the hmmer output and were verified on the CD-search database. Alignments of the BED domains were performed in the same way as for NB-ARC domains and were used to generate a neighbour network in SplitsTree4 based on the uncorrected P distance matrix.

### Transcriptome analysis

#### Kronos analysis

We reanalysed RNA-Seq data from cultivar Kronos^53^ to determine whether the Kronos *Yr5* allele was expressed. We followed the same strategy as that described to define the *Yr7* and *Yr5* gene structures (candidate contig confirmation and gene annotation section). We generated a *de novo* assembly of the Kronos NLR repertoire from Kronos RenSeq data and used it as a reference when mapping read data from one replicate of the wild-type Kronos at heading stage. Read depths up to 30x were present for the *Yr5* allele which allowed confirmation of its expression. Likewise, the RNA-Seq reads confirmed the gene structure, which is similar to *YrSP*, and the premature termination codon in Kronos *Yr5*. Whether this allele confers resistance against *Pst* remains to be elucidated.

#### Re-analysis of RNA-Seq data in Dobon et al., 2016

We used RNA-Seq data previously published by Dobon and colleagues^18^. Briefly, two RNA-Seq time-courses were used based on samples taken from leaves at 0, 1, 2, 3, 5, 7, 9, and 11 dpi for the susceptible cultivar Vuka and 0, 1, 2, 3, and 5 dpi for the resistant AvocetS-Yr5^29^. We used normalised read counts (Transcript Per Million, TPM) from Ramirez-Gonzalez et al. 2018 to produce the heatmap shown in Supplementary Figure 10 with the pheatmap R package^54^ (v1.0.8). Transcripts were clustered according to their expression profile as defined by a Euclidean distance matrix and hierarchical clustering. Transcripts were considered expressed if their average TPM was ≥0.5 TPM in at least one time point. We used the DESeq2 R package^55^ (v1.18.1) to conduct a differential expression analysis. We performed two comparisons: (1) we used a likelihood ratio test to compare the full model ~ Cultivar + Time + Cultivar:Time to the reduced model ~ Cultivar + Time to identify genes that were differentially expressed between the two cultivars at a given time point after 0 dpi (workflow: https://www.bioconductor.org/help/workflows/rnaseq_Gene/); (2) Investigation of both time courses in Vuka and AvocetS-*Yr5* independently to generate all of the comparisons between 0 dpi and any given time point, following the standard DESeq2 pipeline. Genes were considered as differentially expressed genes if they showed an adjusted p-value < 0.05 and a log2 fold change of 2 or higher. Most BED-containing proteins and BED-NLRs were not expressed in the analysed data. No pattern was observed for those that were expressed: differences were observed between cultivars, but these were independent of the presence of the yellow rust pathogen.

## Author contributions

CM performed the experiments to clone *Yr7* and *Yr5* and the subsequent analyses of their loci and BED domains, designed the gene-specific markers, analysed the genotype data in the studied panels, and designed and made the figures. JZ performed the experiments to clone *YrSP*, confirm the *Yr7* and *Yr5* genes in AvocetS-*Yr7* and AvocetS-*Yr5* mutants, and identified the full length of *Yr5* and *YrSP* with their respective regulatory elements. CM and JZ developed the gene specific markers. PZ and RM performed the EMS treatment, isolation, and confirmation of *Yr7, Yr5*, and *YrSP* mutants in AvocetS NILs. PF performed the pathology work on the Cadenza *Yr7* mutants and the mapping populations. BS helped with the NLR annotator analysis and provided the bait library for target enrichment and sequencing of NLRs, NMA provided DNA samples for allelic variation studies and LB provided Lemhi-*Yr5* mutants. RM, EL, PZ, BW, SB, and CU conceived, designed, and supervised the research. CM and CU wrote the manuscript. JZ, PZ, RM, BW, NMA, LB and EL provided edits.

## Data availability

All sequencing data has been deposited in the NCBI Short Reads Archive under accession numbers listed in Supplementary Table 13 (SRP139043). Cadenza (*Yr7*) and Lemhi (*Yr5*) mutants are available through the JIC Germplasm Resource Unit (www.seedstor.ac.uk).

## Acknowledgements

This work was supported by the UK Biotechnology and Biological Sciences Research Council Designing Future Wheat programme BB/P016855/1 and the Grains Research and Development Corporation, Australia. CM was funded by a PhD studentship from Group Limagrain and JZ is funded by PhD scholarships from the National Science Foundation (NSF) and the Monsanto Beachell-Borlaug International Scholars Programs (MBBISP). We thank community feedback on the initial preprint submission which helped improve the manuscript.

We thank the International Wheat Genome Sequencing Consortium for having providing us with pre-publication access to the RefSeq v1.0 assembly and gene annotation. We thank Jorge Dubcovsky and Xiaoqin Zhang (University of California, Davis) for providing *Yr5* cultivars. We thank the John Innes Centre Horticultural Services and Limagrain Rothwell staff for management of the wheat populations. Also Sebastian Specel (Limagrain; Clermont-Ferrand) and Richard Goram (JIC) for their help in designing and running KASP assays, and Sami Hoxha (The University of Sydney) for technical assistance. This research was supported by the NBI Computing Infrastructure for Science (CiS) group in Norwich, UK.

---

**Supplementary Figure 1: Deployment of *Yr7* cultivars in the field is correlated with an increase in the prevalence of *Pst* isolates virulent on *Yr7* in the UK.** Percentage of total harvested weight of wheat cultivar carrying *Yr7* (green) and the proportion of *Pst* isolates that are virulent to Yr7 (orange) from 1990 to 2016 in the United Kingdom. See Supplementary Table 2 for a summary of the data.

**Supplementary Figure 2: Identification of candidate contigs for the *Yr* loci using MutRenSeq.** View of RenSeq reads from the wild-type and EMS-derived mutants mapped to the best candidate contigs identified with MutantHunter for the three genes targeted in this study. From top to bottom: vertical black lines represent the *Yr* loci, coloured rectangles depict the motifs identified by NLR-Annotator (each motif is specific to a conserved NLR domain^56^), while read coverage (grey histograms) is indicated on the left, e.g. [0 - 149], and the line from which the reads are derived on the right, e.g. CadWT for Cadenza wild-type. Vertical bars represent the position of the SNPs identified between the reads and reference assembly – red shows C to T transitions and green G to A transitions. Black boxes highlight SNP for which the coverage was relatively low, but still higher than the 20x detection threshold. The top view shows the *Yr7* allele annotated from the Cadenza genome assembly before manual curation (Supplementary File 3). Vertical black lines illustrate the assembled candidate contigs and the one that was formerly *de novo* assembled from Cadenza RenSeq data, lacking the 5’ region containing the BED domain and thus the Cad903 mutation. The middle view illustrates the *Yr5* locus annotated from the Lemhi-*Yr5 de novo* assembly. The results are similar to those described above for *Yr7*. The full locus was *de novo* assembled. The bottom view illustrates the *YrSP* locus annotated from the AvocetS-YrSP *de novo* assembly with the four identified susceptible mutants all carrying a mutation in the candidate contig. The full locus was *de novo* assembled.

**Supplementary Figure 3: Candidate contigs identified by MutRenSeq are genetically linked to the *Yr* loci mapping interval.** Schematic representation of chromosome 2B from Chinese Spring (RefSeq v1.0) with the positions of published markers linked to the *Yr* loci and surrounding closely linked markers that were used to define their physical position (orange rectangle). The chromosome is depicted as a close-up of the physical locus indicating the positions of KASP markers that were used for genetic mapping (horizontal bars, Supplementary Table 15). Blue colour refers to *Yr7*, red to *Yr5*, and purple to *YrSP*. The black arrow points to the NLR cluster containing the best BLAST hits for *Yr7* and *Yr5/YrSP* on RefSeq v1.0. Coloured lines link the physical map to the corresponding genetic map for each targeted gene (see Methods). Genetic distances are expressed in centiMorgans (cM).

**Supplementary Figure 4: Yr7, Yr5 and YrSP proteins do not encode for a Coiled-Coil domain in the N-terminus.** Graphical outputs from the COILS prediction programm in three sliding windows (14, 21, and 28 amino acid, shown in green, blue, and red, respectively) for Yr5 and Yr7 with or without the BED domain (left) and characterised canonical NLRs: Sr33^21^, Mla10^20^, Pm3^22^ and RPS5^57^. The X axis shows the amino acid positions and the Y axis the probability of a coiled coil domain formation. There was no difference in the prediction between the two Yr proteins with or without their BED domain. The 14 amino acid sliding window is the least accurate according to the user manual, consistent with the additional peaks observed in Sr33, Mla10 and Pm3 that were not annotated as CC domains in the corresponding publications^20–22^. Thus, the peak at position 1,200 in Yr5 is unlikely to represent a CC domain. We performed a BLASTP search with the N-terminal region of the Yr5 and Yr7 proteins (from Met to the first amino-acid encoding the NB-ARC) with or without the BED domain and the best hits were proteins predicted to encode BED-NLRs from *Aegilops tauschii, Triticum uratu* and *Oryza Sativa* (data not shown). Based on the COILS prediction and the BLAST search, we concluded that Yr7 and Yr5/YrSP do not encode CC domains.

**Supplementary Figure 5: Pedigrees of selected Thatcher-derived cultivars and their *Yr7* allelic status.** Pedigree tree of Thatcher-derived cultivars where each circle represents a cultivar and the size of the circle is proportional to its prevalence in the tree. Colours illustrate the genotype with red showing the absence of *Yr7* and yellow its presence. Cultivars in grey were not tested or are intermediate crosses. *Yr7* originated from *Triticum durum* cv. Iumillo and was introgressed into hexaploid wheat through Thatcher (indicated by arrow). Each *Yr7* positive cultivar is related to a parent that was also positive for *Yr7*. Figure was generated using the Helium software^45^ (v1.17).

**Supplementary Figure 6: Diagnostic genetic marker for *Yr5*.** The Yr5-specific insertion was used to generate a PCR amplification product of 1,281 bp for *Yr5* or a shorter amplicon for the absence of the insertion in *YrSP*, Claire, and Paragon (507 bp). *Yr5* positive lines include the *Yr5* spelt donor and *Yr5* near-isogenic lines AvocetS-*Yr5* and Lemhi-*Yr5. YrSP* donor Spaldings Prolific and *YrSP* near-isogenic lines AvocetS-*YrSP* carry the shorter alternate allele, similar to the Claire, Cadenza and Paragon alleles identified in Figure 2. Negative controls include AvocetS and H2O. Size marker is shown on the left.

**Supplementary Figure 7: Expansion of BED-NLRs in the Triticeae and presence of conserved BED-BED-NLRs aross the syntenic region.** Schematic representation of the physical loci containing *Yr7* and *Yr5/YrSP* homologs on RefSeq v1.0 and its syntenic regions. The syntenic region is flanked by conserved non-NLR genes (orange arrows). Black arrows represent canonical NLRs and purple/blue/red arrows represent different types of BED-NLRs based on their BED domain and their relationship identified in Figure 3 and Supplementary Figure 8. Black lines represent phylogenetically related single NLRs located between the two NLR clusters illustrated in Supplementary Figure 9. Details of genes are reported in Supplementary File 4.

**Supplementary Figure 8: The *Yr* loci are phylogenetically related to nearby NLRs on RefSeq v1.0 and their orthologs.** Phylogenetic tree based on translated NB-ARC domains from NLR-Annotator. Node labels represent bootstrap values for 1,000 replicates. The tree was rooted at mid-point and visualized with Dendroscope v3.5.9. The colour pattern matches that of Figure 3 to highlight BED-NLRs with different BED domains.

**Supplementary Figure 9: Neighbour-net analysis network as shown in Figure 3 with identifiers.** Neighbour-net analysis based on uncorrected *P* distances obtained from alignment of 153 BED domains including the 108 BED-containing proteins (including 25 NLRs) from RefSeq v1.0, BED domains from NLRs located in the syntenic region as defined in Supplementary Figure 7, and BED domains from Xa1 and ZBED from rice. BED-I and II clades are highlighted in purple and blue, respectively. BED domains from the syntenic regions not related to either of these types are in red. BED domains derived from non-NLR proteins are in black and BED domains from BED-NLRs outside the syntenic region are in grey. Seven BED domains from non-NLR proteins were close to BED domains from BED-NLRs.

**Supplementary Figure 10: BED-NLRs and BED-containing proteins are not differentially expressed in yellow rust-infected susceptible and resistant cultivars.** Heatmap representing the normalised read counts (Transcript Per Million, TPM) from the reanalysis of published RNAseq data^29^ for all the BED-containing proteins, BED-NLRs and canonical NLRs located in the syntenic region annotated on RefSeq v1.0. Lack of expression is shown in white and expression levels increase from blue to red. Asterisks show cases where several gene models were overlapping with NLR loci identified with NLR Annotator. The colour pattern matches that of Figure 3 to highlight BED-NLRs with different BED domains. Orange labels show the expression of the canonical NLRs located within the syntenic interval. The seven non-NLR BED genes whose BED domain clustered with the ones from BED-NLR proteins in Figure 3 and Supplementary Figure 9 are indicated by black triangles.

**Supplementary Table S1: Summary of *Pst* isolates tested on *Yr5* differential lines from 2004 to 2017 in different regions.** Overall, >6,000 isolates from 44 countries displaying >200 different pathotypes were tested on *Yr5* materials and no virulence was recorded apart from one isolate from Australia, PST 360 E137 A+^14^. Data were obtained from public databases and reports on yellow rust surveillance, whose references are recorded. It is important to note that we report here the number of identified pathotypes for a given region and database. Similar pathotypes could thus have been counted twice if identified in different regions.

**Supplementary Table S2: Harvested weight of known *Yr7* cultivarsfrom 1990 to 2016 and *virYr7* prevalence among UK *Pst* isolates.** Proportion of harvested *Yr7* wheat cultivars in the UK from 1990 to 2016. The prevalence of yellow rust isolates virulent to *Yr7* across this time period is shown in the top row. Original data from NIAB-TAG Seedstats journal (NIAB-TAG Network) and the UK Cereal Pathogen Virulence Survey (http://www.niab.com/pages/id/316/UKCPVS).

**Supplementary Table S3: Plant materials analysed for the present study with the different *Pst* isolates used for the pathology assays.**

**Supplementary Table S4: Plant material submitted for Resistance gene enrichment Sequencing (RenSeq).** From left to right: Mutant line identifier, targeted gene, score when infected with *Pst* according to the Grassner and Straib scale, mutation position, coverage of the mutation (at least 99% of the reads supported the mutant base in the mutant reads), predicted effect of the mutation on the protein sequence, comments. Lines with the same mutations are highlighted with matching colours.

**Supplementary Table S5: Genome assemblies used in the present study.** Summary of the available genome assemblies^58,59^ that were used for the *in silico* allele mining and synteny analysis across rice, *Brachypodium*, barley and different Triticeae accessions.

**Supplementary Table S6: *In silico* allele mining for *Yr7* and *Yr5/YrSP* in available genome assemblies for wheat.** Table presents the percentage identity (% ID) of the identified alleles and matching colours illustrate identical haplotypes. Investigated genome assemblies are shown in Supplementary Table 5.

**Supplementary Table S7: Polymorphisms between Yr5 protein and its identified alleles.** Positions of the polymorphic amino acids across the five Yr5/YrSP proteins. Polymorphisms falling into the BED and NB-ARC domains are shown in red and blue, respectively.

**Supplementary Table S8: Presence/absence of *Yr7* alleles in a selected panel of Cadenza-derivatives and associated responses to different *Pst* isolates (avirulent to *Yr7: Pst* 15/151 and 08/21; virulent to *Yr7:* 14/106).** Infection types were grouped into two categories: 1 for resistant and 2 for susceptible. We used Vuka as a positive control for inoculation and absence of *Yr7*. The typical response of a *Yr7* carrier would thus be 1 – 1 – 2, although some cultivars might carry other resistance genes that can lead to a 1 – 1 – 1 profile (e.g. Cadenza). Cultivars that were positive for *Yr7* had either one or the other profile so none of them was susceptible to a *Pst* isolate that is avirulent to *Yr7*. Few cultivars (e.g Bennington, KWS-Kerrin, Brando) were susceptible to one of the two isolates avirulent to *Yr7* in addition to their susceptibility to the *Yr7*-virulent isolate. However, none of them carried the *Yr7* allele.

**Supplementary Table S9: Presence/absence of *Yr7* and *YrSP* in different wheat collections.** We used Vuka, AvocetS and Solstice as negative controls for the presence of *Yr7* and *YrSP* and AvocetS-Yr near-isogenic lines as controls for the corresponding *Yr* gene. We genotypied different collections: (i) a set of potential *Yr7* carriers based on literature research, (ii) a set of cultivars that belonged to the UK AHDB Recommended List (https://cereals.ahdb.org.uk/varieties/ahdb-recommended-lists.aspx) between 2005 and 2018 (labelled 2005–2018-UK_RL), (iii) the Gediflux collection that includes modern European bread wheat cultivars (1920–2010)^25^, (iv) a core set of the Watkins collection, which represent a set of global bread wheat landraces collected in the 1920-30s^24^. Most of the putative *Yr7* carriers were positive for all the *Yr7* markers apart from Aztec, Chablis and Cranbrook. Chablis was susceptible to the *Pst* isolates that were avirulent to *Yr7* so it probably does not carry the gene. Regarding the 2005-2018-UK_RL results were consistent across already tested cultivars: Cadenza, Cordiale, Cubanita, Grafton and Skyfall were already positive in Supplementary Table 8. Energise, Freiston, Gallant, Oakley and Revelation were negative on both panels as well. Results were thus consistent across different sources of DNA. Yr7-containing cultivars are not prevalent in the 2005-2018 Recommended List set, however, this gene is present in Skyfall, which is currently one of the most harvested cultivars in the UK (Supplementary Table 2>). We tested the *YrSP* marker on this set and it was positive only for AvocetS-*YrSP*. The frequency of *Yr7* was relatively low in the Gediflux panel (4%). This is consistent with results in Supplementary Table 2: *Yr7* deployment started in the UK in 1992 with Cadenza and it was rarely used prior to that date. The same was observed in the subset of the Watkins collection (10%) where landraces that were positive for *Yr7* all originated from India and the Mediterranean basin. *Yr7* was introgressed into Thatcher (released in 1936) from Iumillo, which originated from Spain and North-Africa (Genetic Resources Information System for Wheat and Tritical - http://www.wheatpedigree.net/). Iumillo is likely to be pre-1920s and these landraces are all bread wheats so they might have inherited it from another source. However, there is no evidence for *Yr7* coming from another source than Iumillo in the modern bread wheat cultivars.

**Supplementary Table S10: Presence/absence of *Yr5* alleles in selected cultivars.** A subset of the aforementioned collection was investigated for the *Yr5* presence. “Yes” in the *Yr5* column refers to amplification of the 1,281 bp amplicon with the Yr5-Insertion primers (Supplementary Figure 6). “Yes” in the *Yr5* alternate alleles column refers to the amplification of the 507 bp amplicon that was identified for *AvocetS-YrSP*, Claire, Cadenza and Paragon in Supplementary Figure 6. “Yes” in the no amplification column refers to identification of a profile similar to the one found for AvocetS in Supplementary Figure 6.

**Supplementary Table S11: Identified BED-containing proteins in RefSeq v1.0 based on a hmmer scan analysis (see Methods).** Several features are added: number of identified BED domains and the presence of other conserved domains present, the best BLAST hit from the non-redundant database of NCBI with its description and score, and whether the BED domain was related to BED domains from NLR proteins based on the neighbour network shown in Supplementary Figure 8.

**Supplementary Table S12: Transcripts per Million-normalised read counts from the re-analysis of published RNA-Seq data^29^ and associated differential expression analysis performed with DESeq2.**

**Supplementary Table S13: Sequencing details of RenSeq data generated in this study.**

**Supplementary Table S14: *De novo* assemblies generated from the corresponding RenSeq data.**

**Supplementary Table S15: Primers designed to map and clone *Yr7, Yr5*, and *YrSP*.** Note that KASP assays require the addition of the corresponding 5’-tails for the two KASP primers

**Supplementary Table S16: Diagnostic markers for *Yr7, Yr5*, and *YrSP*.** Note that KASP assays require the addition of the corresponding 5’ -tails for the two KASP primers.

**Supplementary Table S17: Passport data of tested *T. dicoccoides* accessions**

**Supplementary File 1: Annotation of the *Yr7* locus in Cadenza with exon/intron structure, positions of mutations and the position of primers for long-range PCR and nested PCRs that were carried out prior to Sanger sequencing (Supplementary Table 15)**.

The file also includes the derived CDS and protein sequences with annotated conserved domains. Amino acids encoding the BED domain are shown in red and those encoding the NB-ARC domain are in blue. LRR repeats identified with NLR Annotator are highlighted in dark green and manually annotated LRR motifs xxLxLxx are underlined and in bold black.

**Supplementary File 2: Annotation of the *Yr5/YrSP* locus in Lemhi-*Yr5* and AvocetS-*YrSP*, respectively, with exon/intron structure, the position of mutations and the position of primers for long-range PCR and nested PCRs that were carried out prior to Sanger sequencing (Supplementary Table 15)**.

The derived CDS and protein sequences with annotated conserved domains are also shown. Amino acids encoding the BED domain are shown in red and those encoding the NB-ARC domain are in blue. LRR repeats identified with NLR Annotator are highlighted in dark green and manually annotated LRR motifs xxLxLxx are underlined and in bold black. Design of the *Yr5* PCR marker is shown at the end of the file with the insertion that is specific to *Yr5* when compared to *YrSP* and Claire.

**Supplementary File 3: Curation of the *Yr7* locus in the Cadenza genome assembly based on Sanger sequencing results**.

Comments show the position of the unknown bases (“N”) in the “Yr7_with_Ns” sequence. Curation based on Sanger sequencing data is shown in bold black in the “curated_Yr7” sequence with the 39 bp insertion and 129 bp deletion. Allele mining for *Yr7* in the Paragon assembly showed that a similar assembly issue might have occurred for this cultivar (same annotation in the “Yr7_Paragon_with_Ns” sequence). This is consistent with the fact that both assemblies were produced with the same pipeline (Supplementary Table 5). We used RenSeq data available for Paragon and performed an alignment as described for the MutRenSeq pipeline against Cadenza NLRs with the curated *Yr7* loci included. A screen capture of the mapping is shown. Only one SNP was identified (75% Cadenza, 25% Paragon). Across the six reads supporting the alternate base, four displayed several SNPs and mapped to an additional Cadenza NLR. This provides evidence for the presence of the identical gene in Paragon which is supported by phenotypic data.

**Supplementary File 4: Syntenic region across different grasses (Supplementary Table 5) and the NLR loci identified with NLR-Annotator**.

See Methods for a detailed explanation of the analysis and Supplementary Figure 7 for an illustration.

**Supplementary File 5: Curated sequences of BED-NLRs from chromosome 2B and Ta_2D7**.

Exons are highlighted with different colours (yellow, green, blue, pink). Amino acids encoding the BED domain are shown in red and those encoding the NB-ARC domain are in blue. LRR repeats identified with NLR Annotator are highlighted in dark green and manually annotated LRR motifs xxLxLxx are underlined and in bold black.

